# Temporal expectations modulate face image repetition suppression as indexed by event-related potentials

**DOI:** 10.1101/361170

**Authors:** Daniel Feuerriegel, Owen Churches, Scott Coussens, Hannah A.D. Keage

## Abstract

Repeated exposure to a stimulus leads to reduced responses of stimulus-selective sensory neurons, an effect known as repetition suppression or stimulus-specific adaptation. Several influential models have been proposed to explain repetition suppression within hierarchically-organised sensory systems, with each specifying different mechanisms underlying repetition effects. We manipulated temporal expectations within a face repetition experiment to test a critical prediction of the predictive coding model of repetition suppression: that repetition effects will be larger following stimuli that appear at expected times compared to stimuli that appear at unexpected times. We recorded event-related potentials from 18 participants and mapped the spatiotemporal progression of repetition effects using mass univariate analyses. We then assessed whether the magnitudes of observed face image repetition effects were influenced by temporal expectations. In each trial participants saw an adapter face, followed by a 500ms or 1000ms interstimulus interval (ISI), and then a test face, which was the same or a different face identity to the adapter. Participants’ expectations for whether the test face would appear after a 500ms ISI were cued by the sex of the adapter face. Our analyses revealed multiple repetition effects with distinct scalp topographies, extending until at least 800ms from stimulus onset. An early (158-203ms) repetition effect was larger for stimuli following surprising, rather than expected, 500ms ISI durations, contrary to the model predictions of the predictive coding model of repetition suppression. Later (230-609ms) repetition effects tended to be larger following expected stimulus onset times, in line with predictive coding models. Our results indicate that the relationship between repetition suppression and temporal expectation differs across the time course of the stimulus-evoked response, suggesting multiple distinct mechanisms driving repetition suppression that operate at different latencies within the visual hierarchy.

**Highlights:** - Multiple face image repetition effects identified from 162-800ms post stimulus onset

- Temporal expectations influenced the magnitudes of repetition effects

- Temporal expectation effects differed for early and late stimulus-evoked responses

## 1. Introduction

Living organisms exhibit a remarkable ability to exploit statistical regularities and recurring patterns that occur within sensory environments. Many vertebrate and invertebrate species can rapidly form predictions based on recurring sequences of stimuli, allowing them to anticipate the identities, locations and timing of upcoming events (e.g., Posner, 1980; Turk-Browne et al., 2009; Meyer & Olson, 2011; Hogendoorn & Burkitt, 2018; Nobre & van Ede, 2018). Responses of sensory neurons are also shaped by recent stimulus exposure. Repeated presentation of a stimulus typically leads to reductions in responses of stimulus-selective cortical and subcortical neurons, known as repetition suppression or stimulus-specific adaptation (Desimone, 1996; Movshon & Lennie, 1979).

Repetition suppression (RS) refers to a stimulus-specific reduction in a recorded signal of neuronal activity (e.g. firing rate, local field potential amplitude, fMRI BOLD signal change) to repeated compared to unrepeated stimuli (Henson et al., 2004; De Baene & Vogels, 2010; for reviews see Grill-Spector et al., 2006; Kohn, 2007; Vogels, 2016; Larsson et al., 2016). Repetition effects have also been reported when recording EEG/MEG (e.g., Caharel et al., 2015; Feuerriegel et al., 2018a). These effects are widely believed to index RS, due to almost ubiquitous findings of suppression (rather than enhancement) of neural responses when using similar experimental designs combined with different recording modalities (e.g., single unit firing rates: Sawamura et al., 2006; local field potentials: De Baene & Vogels, 2010; fMRI BOLD signals: Grill-Spector et al., 1999). Characterising the neural mechanisms that underlie RS is critical for understanding how we detect rare or novel events in our environment (Nelken, 2014; Solomon & Kohn, 2014), and what occurs when this process functions abnormally in neurological and psychiatric disorders (e.g., Naatanen et al., 2014, Kremlacek et al., 2016). Here we will focus on *immediate* stimulus repetition, as opposed to *delayed* repetition (i.e., when several intervening stimuli are presented between the first and repeated presentations of a stimulus – see Henson, 2016).

Several conceptual and computational models have been proposed to explain RS. Early models described local mechanisms that influence the rate, duration, and stimulus selectivity of neural responses (Desimone, 1996; Wiggs & Martin, 1998; reviewed in Grill-Spector et al., 2006). More recent models acknowledge that RS operates within hierarchically organised sensory systems, such as the visual system. These newer models emphasize that repetition effects occur within local, recurrently-connected neural networks, and can be propagated across brain regions. There are currently two dominant models of RS, which both focus on response modulations of stimulus-selective excitatory neurons, such as cortical pyramidal neurons which contribute to scalp-recorded EEG.

Normalisation models (Dhruv et al., 2011; Solomon & Kohn, 2014; Kaliukhovich & Vogels, 2016; Whitmire & Stanley, 2016) describe responses of stimulus-selective neurons according to the interplay between excitatory (i.e., driving afferent) input, corresponding to stimulation within classical receptive fields, and divisive normalising inhibitory input from other neurons within the same network. These are similar networks to those specified in normalisation models of attention (e.g., Reynolds & Heeger, 2009). The nature of the divisive normalising input differs by brain region (reviewed in Carrandini & Heeger, 2012), for example expressed as inhibitory ‘surround’ effects in V1 (Wissig & Kohn, 2012) or competitive interactions between feature-selective neurons in extrastriate visual areas, possibly acting via GABAergic interneurons (e.g., Chelazzi et al., 1998; Kaliukhovich & Vogels, 2016). Importantly, the effects of both excitatory and inhibitory inputs can be reduced by stimulus exposure in these models, for example due to spike frequency adaptation, afterhyperpolarisation or rapid synaptic plasticity (Zucker & Regehr, 2002; Fioravante & Regehr, 2011; reviewed in Whitmire & Stanley, 2016; Vogels, 2016). RS can also propagate across visual areas in a feedforward or feedback manner, due to downstream visual areas receiving altered input from adapted neural populations (e.g., Kohn, & Movshon, 2003; Dhruv & Carrandini, 2014). Such models postulate that a primary function of RS is to increase the salience of novel stimuli by enhancing responses to novel stimuli compared to those seen in the recent past.

Another dominant model of RS is derived from theories of perception based on predictive coding (e.g., Rao & Ballard, 1999; Friston, 2005) and is described in detail in Auksztulewicz and Friston (2016). Predictive coding models describe RS as a reduction of prediction error signals, due to fulfilled perceptual expectations that are weighted toward recently-encountered stimuli. In this model reductions in responses of superficial pyramidal neurons (which signal prediction errors) occur via inhibitory lateral and feedback connections (Friston, 2005), for example via GABAergic inhibitory interneurons (Chu et al., 2003; Wozny & Williams, 2011). A critical component of this model is sensory precision, which reflects the confidence that a system holds regarding its sensory predictions (Feldman & Friston, 2010). Accordingly, prediction errors are weighted by the sensory precision of predictions. Sensory precision can be manipulated by exogenous factors, such as stimulus signal-to-noise ratio, or endogenous factors, such as focused attention, or the expectation that a certain stimulus will appear (Feldman & Friston, 2010; Auksztulewicz & Friston, 2016). According to the predictive coding model of RS, contexts associated with high sensory precision (i.e., attended and/or expected stimuli) are predicted to lead to larger RS, compared with contexts associated with lower precision (i.e., unattended and/or surprising stimuli).

To test and extend normalisation and predictive coding models of RS researchers have assessed whether RS is modulated by attention and expectation. Studies manipulating attention within immediate repetition designs have reported larger fMRI BOLD RS for stimuli of attended (compared to unattended) spatial locations and stimulus categories (Murray & Wojciulik, 2004; Eger et al., 2004; Yi et al., 2006). The N250r ERP face identity repetition effect was also reduced when attention was diverted towards a distractor face (Neumann & Schweinberger, 2009). These effects of attention are congruent with predictive coding accounts of RS, and signify that normalisation models could be extended to describe interactions between attention and RS.

There is also a substantial literature on RS and perceptual expectations (e.g., expectations based on the contextual likelihood that a given stimulus will appear). Summerfield and colleagues (2008) presented pairs of repeated (i.e., AA) or alternating (i.e., AB) faces in each trial, and manipulated across blocks the proportions of trials with face repetitions (60% vs. 20%). They reported larger face identity BOLD RS in the fusiform face area (FFA; Kanwisher et al., 1997) in blocks with higher proportions of repetition trials compared to those with lower proportions. These findings have been replicated several times using fMRI (e.g., Kovács et al., 2012; 2013; de Gardelle et al., 2013; Grotheer & Kovács, 2014; Choi et al, 2017), and have been widely interpreted as increased RS resulting from higher sensory precision in contexts whereby repetitions were expected to occur. However, the analyses used in these experiments confounded additive and interactive effects of RS and expectation (discussed in Grotheer & Kovács, 2015; Feuerriegel et al., 2018a). Other experiments independently manipulated stimulus repetition and expectation using fMRI and electrophysiological recordings, and observed independent expectation and RS effects (Todorovic & de Lange, 2012; Kaliukhovich & Vogels, 2011; 2014; Grotheer & Kovács, 2015; Feuerriegel et al., 2018a). Another study instead observed larger RS for surprising, rather than expected, stimuli (Amado et al., 2016); this pattern of effects is also visible in many of the Summerfield et al. replications (Kovács et al., 2012; de Gardelle et al., 2013; Larsson & Smith, 2012; Grotheer & Kovács, 2014; Choi et al., 2017; reviewed in Kovács & Vogels, 2014). Findings of larger RS for surprising (rather than expected) stimuli are the opposite pattern to that hypothesized by the predictive coding model (Auksztulewicz & Friston, 2016).

In the current study we designed a different test of the precision-based modulation hypothesis; we manipulated expectations regarding *when* an upcoming stimulus would appear (i.e., temporal expectations, Nobre et al., 2007; Nobre & van Ede, 2018). There appear to be multiple, widespread effects of temporal expectation in the brain (reviewed in Nobre & van Ede, 2018). In primate visual cortex temporal expectations can increase firing rates and local field potential amplitudes to expected stimuli in V4 and inferior temporal cortex (Ghose & Maunsell, 2002; Anderson & Sheinberg, 2007), and drive increased gamma band oscillations and suppressed alpha-band activity in V1 (Lima et al., 2011), similar to effects of spatial attention (Fries et al., 2008). Temporal expectations can also modulate visual stimulus evoked potentials and amplify the effects of spatial attention on scalp-recorded ERPs (Doherty et al., 2005; Correa et al., 2006). According to predictive coding models of RS, stimuli that appear at expected times are linked to higher sensory precision, and such stimuli should show larger RS than those which appear at unexpected/surprising times (Auksztulewicz & Friston, 2016).

It is currently unclear whether temporal expectations influence stimulus repetition effects in the visual system. Differences in ERP repetition effect magnitudes have been reported for auditory stimuli in oddball designs, which presented streams of stimuli separated by isochronous or random interstimulus intervals (Costa-Faidella et al., 2011; Schwartze et al., 2013; but see experiments 1 and 2 in Tavano et al., 2014). In these experiments repetition effects were operationalised as the additive effects of stimulus repetition and stimulus feature expectations; repeated stimulus tones were expected, whereas unrepeated tones were surprising. Consequently, it is unclear whether temporal expectations modulated RS-specific processes or effects of stimulus feature expectations (e.g. Tavano et al., 2014), which would lead to similar patterns of effects on recorded ERPs.

To provide a more specific test of temporal expectation effects on RS, we presented pairs of repeated and alternating (unrepeated) faces separated by 500ms and 1000ms interstimulus intervals (ISIs). We adapted the design of Grotheer and Kovács (2015) to cue participants’ expectations for a 500ms or 1000ms ISI, depending on the sex of the first face presented in each trial. Unlike previous studies we also balanced expectations for specific stimulus identities, and temporal expectations, across repeated and alternating stimuli. By recording ERPs evoked by repeated and alternating faces we tested whether the N250r repetition effect, which is influenced by feature-based attention (Neumann & Schweinberger, 2009), could also be modulated by temporal expectations. Using mass univariate analyses we could also map the complex spatiotemporal progression of repetition effects (e.g., Feuerriegel et al., 2018a), and test whether earlier or later effects are modulated by temporal expectations. Predictive coding models (e.g. Auksztulewicz & Friston, 2016) hypothesise larger repetition effects for stimuli with expected onset times due to increased precision of sensory predictions.

## 2. Methods

### 2.1 Participants

Eighteen people (4 males) participated in this experiment (age range 18-32 years, mean age 23.6 ± 4.9). All participants were native English speakers and had normal or corrected-to-normal vision, no history of psychiatric or neurological disorders or substance abuse, no history of unconsciousness for greater than 1 minute, and had not taken recreational drugs within the last 6 months. All participants were right-handed as assessed by the Flinders Handedness Survey (Nicholls et al., 2013). This study was approved by the Human Research Ethics committee of the University of South Australia.

### 2.2 Stimuli

We took 49 frontal images of faces (24 male, 25 female) from the Karolinska Directed Emotional Faces database (Lundqvist et al., 1998). Examples of stimuli are shown in Figure 1A. Selected faces were of neutral expression with no facial piercings or hair that occluded the face. We then converted all images to greyscale, and cropped, resized and aligned them, so that the nose was in the horizontal center of the image. We then vertically aligned the eyes of each face, and resized the images so that at a viewing distance of 60cm stimuli subtended approximately 3.15° × 3.72° of visual angle (134 × 156 pixels). We created test stimuli to be 20% larger than adapter stimuli to minimise low-level or retinal adaptation. We used the SHINE toolbox (Willenbockel et al., 2010) to equate mean pixel intensity, contrast and Fourier amplitude spectra across the images (Mean normalised pixel intensity = 0.52, RMS contrast = 0.16). Stimuli were presented against a grey background (normalised pixel intensity = 0.52).

**Figure 1.**
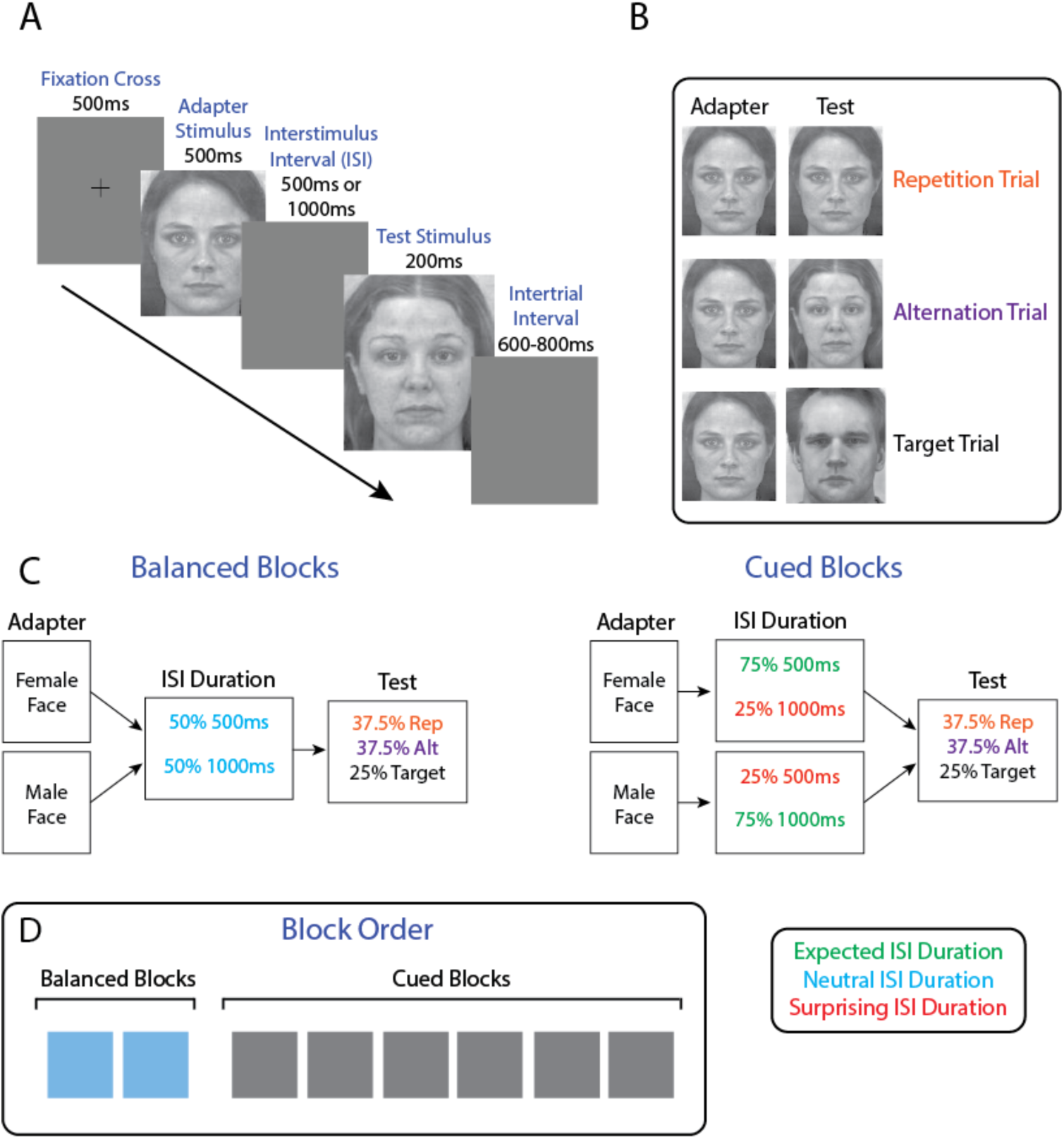
Trial diagram and experimental block types. A) In each trial adapter and test stimuli were presented, separated by either a 500ms or 1000ms ISI. Test stimuli were 20% larger than adapters. An example of an alternating trial is displayed, in which adapter and test faces are different identities. B) Examples of stimuli presented in repetition, alternation and target trials. C) Trial structures for each block type. In balanced blocks the probability of a 500ms or 1000ms ISI duration was 50% each. In cued blocks the probability of a 500ms or 1000ms ISI duration varied by the sex of the adapter face. In this example a female adapter face cues a high (75%) probability of a 500ms ISI, whereas the male adapter face cues a low (25%) probability of a 500ms ISI. Rep and Alt stand for repeated and alternating (i.e., unrepeated) test stimuli. D) Block order in the experiment. Two balanced blocks were presented before 6 cued blocks.

### 2.3 Procedure

Participants sat in a well-lit testing room 60cm in front of an LED monitor (refresh rate 60Hz). We presented stimuli via custom scripts written in MATLAB r2014a (The Mathworks, USA) using functions from PsychToolbox v3.0.11 (Brainard, 1997; Kleiner et al., 2008). Behavioural responses were recorded using a one-button response box.

In each trial faces were presented as adapter stimuli (500ms) and test stimuli (200ms) separated by either a 500ms or 1000ms ISI (Figure 1A). Adapters were preceded by a fixation cross for 500ms. The intertrial interval (including the fixation cross duration) varied pseudorandomly between 1100-1300ms (mean duration = 1200ms). In each block we presented 4 non-target faces (2 male and 2 female) and one target face (either male or female). In non-target trials the adapter stimulus could be any of the four non-target faces, and the test stimulus could either be a repetition of the same face image (repetition trial) or the other face identity of the same gender (alternation trial; see Figure 1B). The proportion of trials with face repetitions stayed constant at 37.5%. To prevent across-trial immediate repetition effects the adapter face in one trial could not be the same identity as the test face in the previous trial. Each non-target face appeared equally as adapters and tests, and as repetition and alternating trial stimuli.

There were eight experimental blocks in the experiment. Within the experiment we manipulated the probability that the test face would appear after a 500ms or 1000ms ISI (see Figure 1C). In the first two blocks the test face appeared after a 500ms or 1000ms ISI with equal (50%) probability (neutral expectation conditions). These were labelled as ‘balanced’ blocks. In the remaining six blocks the probability of a 500ms or 1000ms ISI was cued by the sex of the adapter face. For example, for one participant a female adapter face cued a 75% probability of a 500ms ISI (expected 500ms ISI condition) and a 25% probability of a 1000ms ISI (surprising 1000ms ISI condition), whereas a male adapter face instead cued a 25% probability of a 500ms ISI. These blocks are accordingly labelled as ‘cued’ blocks. Block order if illustrated in Figure 1D. We counterbalanced the adapter face sex used to cue each ISI probability across participants. When questioned after testing no participants reported awareness of this cued ISI probability manipulation.

In target trials (25% of all trials) the target face was presented as the test stimulus. We displayed target faces onscreen during the break before each block for participants to memorise. Target faces allocated to each block were counterbalanced across participants. We instructed participants to press a button with their index finger as quickly as possible after seeing the target face (response hands counterbalanced across participants). We considered responses between 200-1000ms from test stimulus onset as correct responses. The number of male and female target faces was equated within balanced and cued block types. Participants completed a short practice block (24 trials) before the main experiment. We presented a separate set of 5 face images during the practice block, which did not appear in the main experiment.

There were 1600 non-target trials in total: 80 trials for each neutral/balanced (50% probability) and surprising (25% probability) ISI condition, and 240 trials for each expected (75% probability) ISI condition. There were 540 target trials: 54 trials for each neutral and surprising ISI target, and 162 trials for each expected ISI target. Participants were allowed self-paced breaks between blocks. The total time required to complete the experiment (excluding breaks) was 94.5 minutes.

### 2.4 EEG Recording and Data Processing

We recorded EEG from 128 active electrodes using a Biosemi Active Two system (Biosemi, the Netherlands). Recordings were grounded using common mode sense and driven right leg electrodes (http://www.biosemi.com/faq/cms&drl.htm). We added 8 additional channels: two electrodes placed 1cm from the outer canthi of each eye, four electrodes; one placed above and below the centre of each eye, and two electrodes; one placed on each of the left and right mastoids. EEG was sampled at 1024Hz (DC-coupled with an anti-aliasing filter, −3dB at 204Hz). Electrode offsets were kept within ±50*μ*V.

We processed EEG data using EEGLab V.13.4.4b (Delorme and Makeig, 2004) and ERPLab V.4.0.3.1 (Lopez-Calderon and Luck, 2014) running in MATLAB r2015a. We first downsampled EEG data to 512Hz offline. We used a photosensor to measure the timing delay of the video system (10ms) and shifted stimulus event codes offline to account for this delay. We identified 50Hz line noise using Cleanline (Mullen, 2012) using a separate 1Hz high-pass filtered dataset (EEGLab Basic FIR Filter New, zero-phase, finite impulse response, −6dB cutoff frequency 0.5Hz, transition bandwidth 1Hz). We then subtracted the identified line noise from the unfiltered dataset (as recommended by Bigdely-Shamlo et al., 2015). We identified excessively noisy channels by visual inspection (median noisy channels by participant = 1, range 0-4) and excluded these from average referencing and independent components analysis (ICA) procedures. We then re-referenced the data to the average of the 128 scalp channels. We additionally removed one channel (FCz) to correct for the data rank deficiency caused by average referencing. A separate dataset was processed in the same way, except a 1Hz high-pass filter was applied (filter settings as above) to improve stationarity for the ICA. We then performed ICA on the 1Hz high-pass filtered dataset (RunICA extended algorithm, Jung et al., 2000) and transferred the resulting independent component information to the unfiltered dataset. We identified and removed independent components associated with ocular and muscle activity, according to guidelines in Chaumon et al. (2015). Following ICA, we interpolated any noisy channels and FCz using the cleaned data (spherical spline interpolation). We then low-pass filtered the EEG data at 30Hz (EEGLab Basic Finite Impulse Response Filter New, zero-phase, −6dB cutoff frequency 33.75Hz, transition band width 7.5Hz). Data were epoched from −100ms to 800ms from test stimulus onset and baseline-corrected using the prestimulus interval. We excluded from analyses epochs containing ±100μV deviations from baseline, as well as non-target trials containing button press responses.

### 2.5 Statistical Analyses

#### 2.5.1 Behavioural data

We compared mean accuracy percentages and reaction times for targets after expected and surprising 500ms ISIs at the group level, using 20% trimmed means of the within-subject expected/surprise difference scores and 95% confidence intervals derived from the percentile bootstrap method (10,000 bootstrap samples; Efron and Tibshirani, 1993; Wilcox, 2012). For each bootstrap sample the 20% trimmed mean of the difference scores was calculated. From this distribution the values of the 2.5^th^ and 97.5^th^ percentiles were chosen as the edges of the two-tailed 95% confidence interval. This method gives more accurate probability coverage compared to tests based on the arithmetic mean, and is more robust against problems caused by skew and outliers (Wilcox & Keselman, 2003; Wilcox, 2012). Responses to targets following 1000ms ISIs were not compared across expectation conditions, as even when participants expected a 500ms ISI, if the target did not appear by 500ms then it could always be expected to appear after 1000ms (see Nobre et al., 2007).

#### 2.5.2 Mass univariate ERP analyses of stimulus repetition effects

To characterise the spatiotemporal pattern of face image repetition effects we compared ERPs evoked by all repeated and alternating faces (pooled across temporal expectation and ISI conditions) using paired-samples mass-univariate analyses, with cluster-based permutation tests to correct for multiple comparisons, implemented in the LIMO EEG toolbox V1.4 (Pernet et al., 2011). These cluster-based multiple comparisons corrections were used because they provide control over the weak family-wise error rate while maintaining high sensitivity to detect broadly-distributed effects (Maris & Oostenveld, 2007; Groppe et al., 2011). Paired-samples tests were performed at all time points between −100 and 800s at all 128 scalp electrodes (59,008 comparisons) using the paired samples version of Yuen’s t test (Yuen, 1974). Corrections for multiple comparisons were performed using spatiotemporal cluster corrections based on the cluster mass statistic (Bullmore et al., 1999; Maris & Oostenveld, 2007). Paired-samples t tests were performed using the original data and 1000 bootstrap samples. For each bootstrap sample data from both conditions were mean-centred, pooled and then sampled with replacement and randomly allocated to each condition (bootstrap-t method). For each bootstrap sample all t statistics corresponding to uncorrected p-values of <0.05 were formed into clusters with any neighbouring such t statistics. Channels considered spatial neighbours were defined using the 128-channel Biosemi channel neighbourhood matrix in the LIMO EEG toolbox (Pernet et al., 2011; 2015). Adjacent time points were considered temporal neighbours. The sum of the t statistics in each cluster is the ‘mass’ of that cluster. We used the most extreme cluster masses in each of the 1000 bootstrap samples to estimate the distribution of the null hypothesis. We compared the cluster masses of each cluster identified in the original dataset to the null distribution; the percentile ranking of each cluster relative to the null distribution was used to derive its p-value. We assigned the p-value of each cluster to all members of that cluster. Electrode/timepoint combinations not included in any statistically significant cluster were assigned a p-value of 1.

#### 2.5.3 Mass univariate ERP analyses of stimulus repetition effects (localiser dataset)

A second mass univariate analysis was conducted on ERPs to test stimuli in balanced blocks (with 50% probability of a 500ms ISI); this was our localiser dataset to define regions of interest (ROIs) for analysing interactions between stimulus repetition and temporal expectations in the cued blocks. Repeated and alternating stimuli (pooled across 500ms and 1000ms ISIs) were compared using cluster-based permutation tests as described above. We used clusters of statistically-significant repetition effects identified using data from the balanced blocks to define ROIs for testing for expectation by repetition interactions for responses to expected and surprising stimuli in cued blocks.

While others have used mass univariate analyses of repetition effects to derive ROIs for notionally orthogonal expectation x repetition interaction effects within the same dataset (e.g., Summerfield et al., 2011) we instead chose to use an independent localiser dataset. This is because unequal group-level variances across conditions (e.g., when trial numbers are not balanced across expected and surprising conditions) can lead to inflated false positive rates when defining ROIs using orthogonal contrasts (see Brooks et al., 2017; Kriegeskorte et al., 2009).

#### 2.5.4 Repetition effect ROI mean amplitude analyses

For each positive-going and negative-going repetition effect identified using the localiser dataset, we calculated cluster mean ERP amplitudes for repeated and alternating stimuli, following expected and surprising 500ms ISIs. As we were interested in assessing effects of temporal expectations, we did not analyse ERPs evoked by stimuli following 1000ms ISIs in cued blocks. This is because an observer can expect the test stimulus to appear after 1000ms with certainty if it had not already appeared after 500ms had elapsed. Cluster mean amplitudes were calculated as the 20% trimmed mean of all channel/timepoint combinations included within each cluster. Trimmed means were used to minimize effects of skewed distributions and outliers within ROIs, which can influence ROI-averaged measures of neuroimaging data (Friston et al., 2006). Cluster mean amplitudes for each alternating stimulus condition were subtracted from their corresponding expectation-matched repeated stimulus condition, to derive a cluster mean amplitude repetition effect measure. As the positive- and negative-going repetition effects in the localiser dataset appear to be modulations of the same dipolar sources (see Figure 3C) we combined the positive and negative repetition effects into one measure. We did this by subtracting the negative cluster repetition effect mean amplitudes from the positive cluster effect mean amplitudes. This was done to reduce the number of comparisons in these analyses.

Repetition effects for stimuli following expected and surprising 500ms ISIs were then compared using the percentile bootstrap method with 20% trimmed means (Wilcox, 2012). In this analysis framework statistically significant differences in the magnitude of repetition effects for stimuli following expected compared to surprising ISIs are equivalent to a temporal expectation by repetition interaction. The Holm-Bonferroni method (Holm, 1979) was used to correct for multiple comparisons across ROIs. This ROI mean amplitude-based approach allowed us to reduce the number of statistical comparisons as compared to mass univariate analyses, and identify attention or expectation effects that may slightly differ in latency from test stimulus onset across individuals (as done by Summerfield et al., 2011).

In addition to clusters identified using the localiser data, an additional ROI was added, spanning 230-347ms at bilateral occipiotemporal electrodes P7/8, P9/10, PO7/8, and PO9/10, corresponding to the N250r ERP face repetition effect (Schweinberger et al., 2002). This ROI was defined based on the time window during which repeated faces evoked more negative-going waveforms compared to unrepeated faces at these channels in the localiser data grand-averaged ERPs. This N250r effect is a robust face repetition effect, and is of high interest in our study as it was found to be modulated by attention (Neumann & Schweinberger, 2009). The time range and electrodes selected for N250r analyses are consistent with those of previous studies (e.g. Neumann & Schweinberger, 2009).

## 3. Results

### 3.1 Task Performance

We first assessed whether participants were performing the task correctly, and whether the ISI expectation manipulation affected behavioural responses to target trials. Accuracy for detecting and responding to targets, collapsed across conditions, was near ceiling (20% trimmed mean = 99% range 93-100%). Accuracy did not differ for targets following expected compared to surprising ISIs (trimmed mean accuracy scores ranged between 98-100% across conditions). The trimmed mean reaction time to target faces, collapsed across conditions, was 479ms (range 371-642ms across participants). There were no statistically significant differences in reaction times to targets after expected compared to surprising ISIs, for targets after 500ms and 1000ms ISIs (trimmed mean difference for 500ms ISI = −3ms, 95% CI = [−10.7, 3.5], p = .20; difference for 1000ms ISI = 1.6ms, CI = [−6.6, 8.8], p = .34).

### 3.2 Mass Univariate Analyses of Face Image Repetition Effects

#### 3.2.1 Analyses using all non-target trials

To characterise the spatiotemporal pattern of face image repetition effects we conducted mass univariate analyses of ERPs comparing responses to repeated and alternating test stimuli. These analyses revealed four time periods with distinct topographical patterns of stimulus repetition effects (shown in Figure 2A-C). The earliest repetition effect (labelled as Cluster 1) spanned 162-211ms from test stimulus onset, during which repeated stimuli evoked more positive waveforms at bilateral occipitotemporal channels and more negative waveforms at frontocentral channels. A later effect (Cluster 2) spanned 228-369ms, during which waveforms to repeated stimuli were more negative at left occipitotemporal channels and more positive at frontal sites. Cluster 3 spanned 371-619ms and consisted of more negative-going waveforms to repeated stimuli at bilateral posterior sites centred around Pz (but extending to PO7 and PO8), accompanied by more positive-going waveforms at frontal channels. The last cluster (Cluster 4) spanned 720-800ms, and had a similar topography to the first cluster, with more positive-going waveforms at occipitotemporal channels to repeated stimuli, and more negative-going waveforms at central channels.

**Figure 2.**
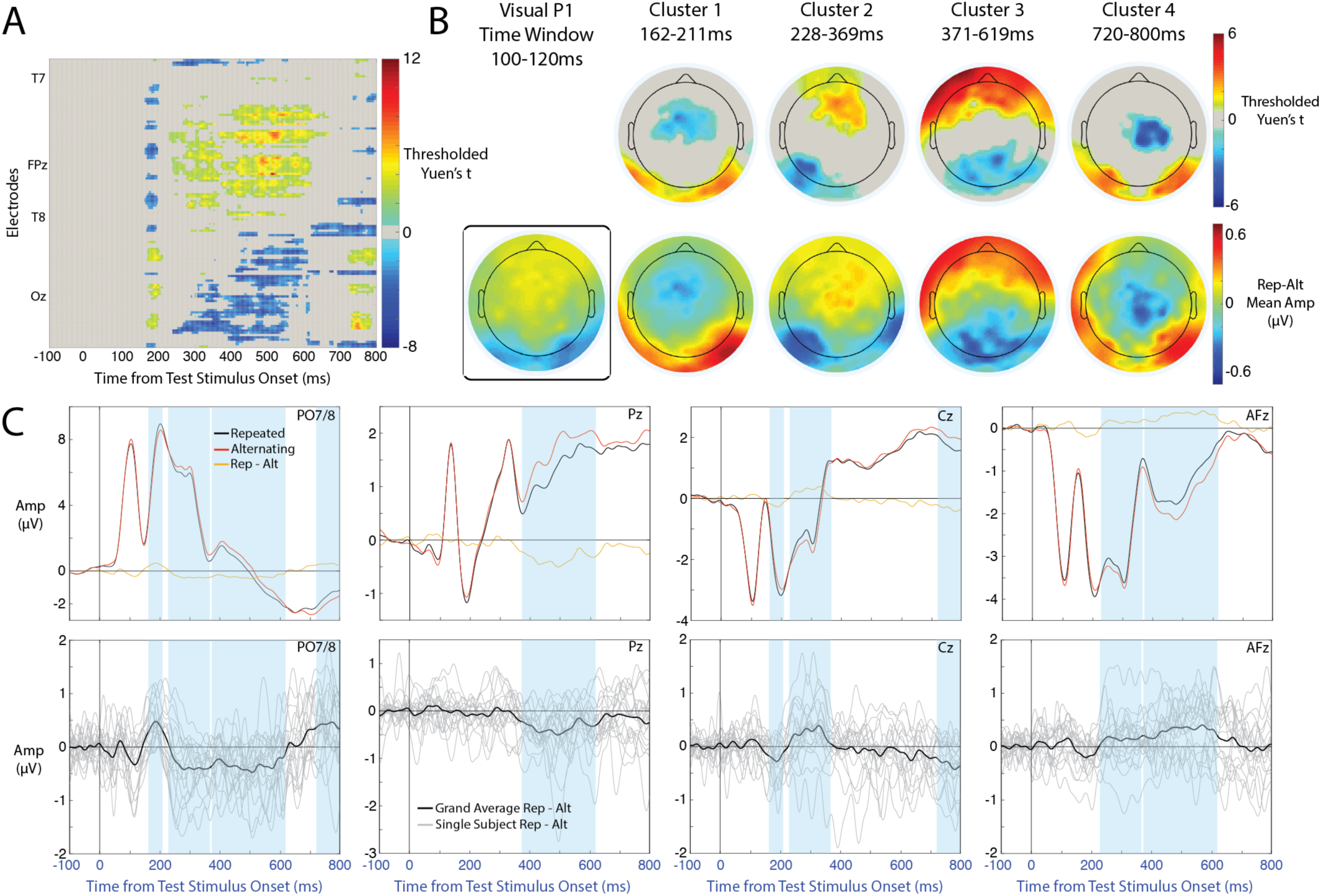
Results of mass univariate analyses of repetition effects. A) Spatiotemporal map of statistically significant repetition effects. Yuen’s t is plotted for each channel/timepoint combination, thresholded by cluster-level statistical significance. B) Scalp maps are displayed for each of 4 identified time windows showing distinct topographies of repetition effects. The top row displays the mean Yuen’s t value over the time window of each cluster. The bottom row shows the [repetition – alternation] average amplitude differences over each time window. The repetition effect during the visual P1 time window was not statistically significant, but was plotted for comparison with Feuerriegel et al. (2018a). C) Grand-averaged ERPs to repeated and alternating stimuli (top row) and grand-average and single-subject repetition-alternation difference waveforms (bottom row). Blue shaded areas denote time windows of statistically significant clusters within which the plotted channel was included.

An earlier repetition effect could also be observed in the group-averaged and single-subject ERPs at electrodes PO7/8, during the time window of the P1 component (100-120ms; Figure 2C). Although this repetition effect was not statistically significant, the topography of this effect (depicted in Figure 2B) was highly similar to that found in our recent study (Feuerriegel et al., 2018a).

#### 3.2.2 Analyses using the localiser dataset

We also ran the same face image repetition effect analyses using data only from the balanced blocks, in order to derive ROIs for analyses of temporal expectation by repetition interactions in the cued blocks. We identified 2 statistically significant clusters (displayed in Figure 3A, 3C). An early repetition effect (Localiser Cluster 1) was observed spanning 158-203ms, during which repeated stimuli evoked more positive waveforms at bilateral occipitotemporal channels and more negative waveforms at frontocentral channels (similar to Cluster 1 in Figure 2B). A later repetition effect (Localiser Cluster 2) spanned 345-609ms and consisted of more negative-going waveforms to repeated stimuli at posterior sites (centred around Pz) accompanied by more positive-going waveforms at frontal channels (similar to Cluster 3 in Figure 2B).

**Figure 3.**
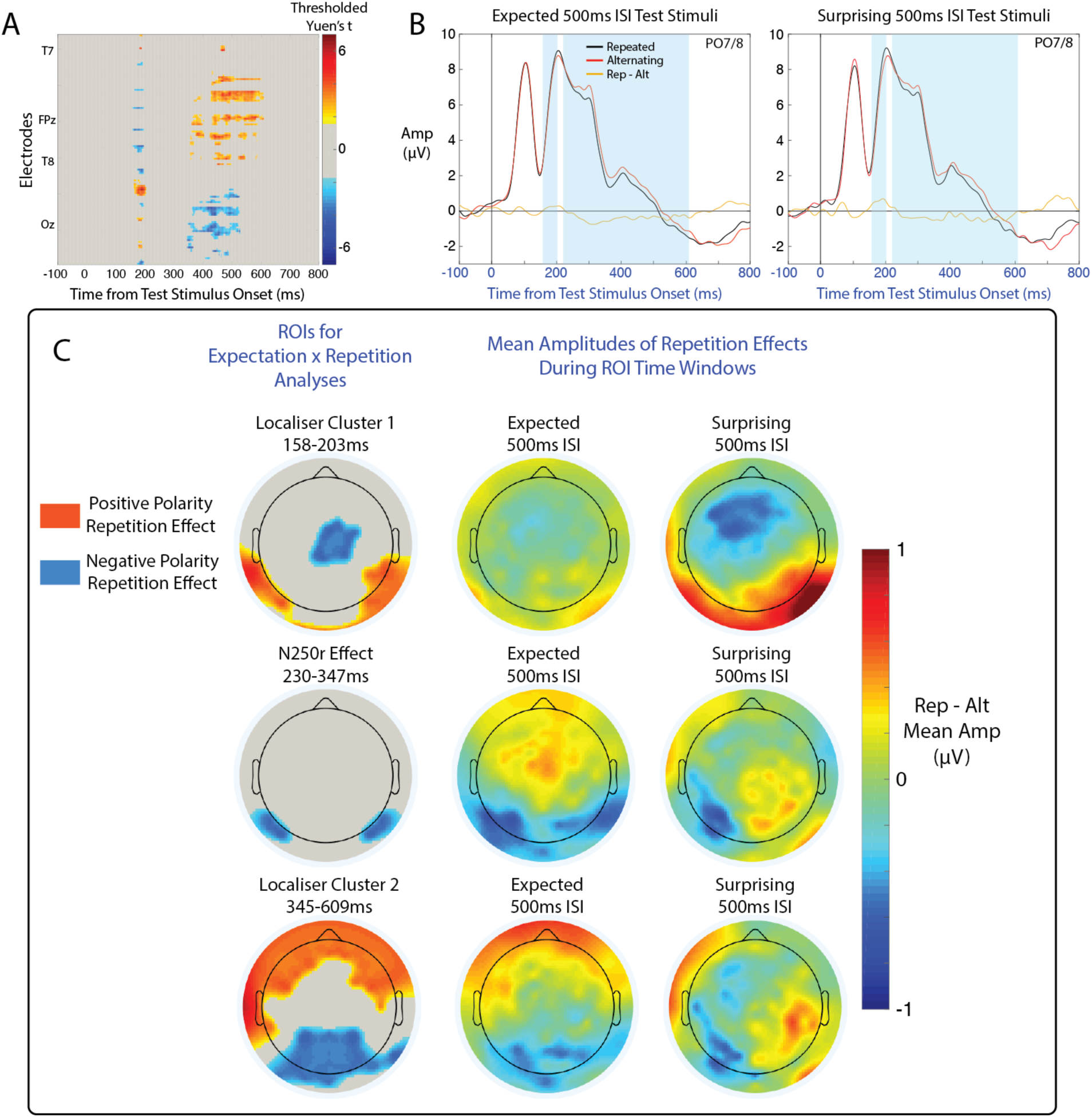
Repetition effect clusters derived from the localiser data, and estimates of repetition effects for stimuli with expected and surprising onset times. A) Repetition effects from mass univariate analyses on the localiser dataset. Yuen’s t is plotted for each channel/timepoint combination, thresholded by cluster-level statistical significance. B) Grand-average ERPs to repeated and alternating stimuli for expected and surprising 500ms ISI conditions. Blue shaded areas denote ROI time windows selected for analysis based on localiser data repetition effects and the N250r ROI definition. C) Topographies of localiser-derived ROIs (left column) and mean amplitudes of repetition effects during ROI time windows for expected and surprising stimuli (center and right columns). In maps of localiser ROIs Electrodes included within a positive repetition effect cluster are coloured red, electrodes included in a negative repetition effect cluster are coloured blue.

### 3.3 ROI Mean Amplitude Analyses

After deriving ROIs using the localiser dataset we then assessed whether repetition effects captured by these ROIs were modulated by temporal expectations. Grand-average ERPs displaying repetition effects for stimuli following expected and surprising 500ms ISIs are displayed in Figure 3B. Estimates of ROI-averaged repetition effects for stimuli following expected and surprising 500ms ISIs are displayed in Figure 4.

**Figure 4.**
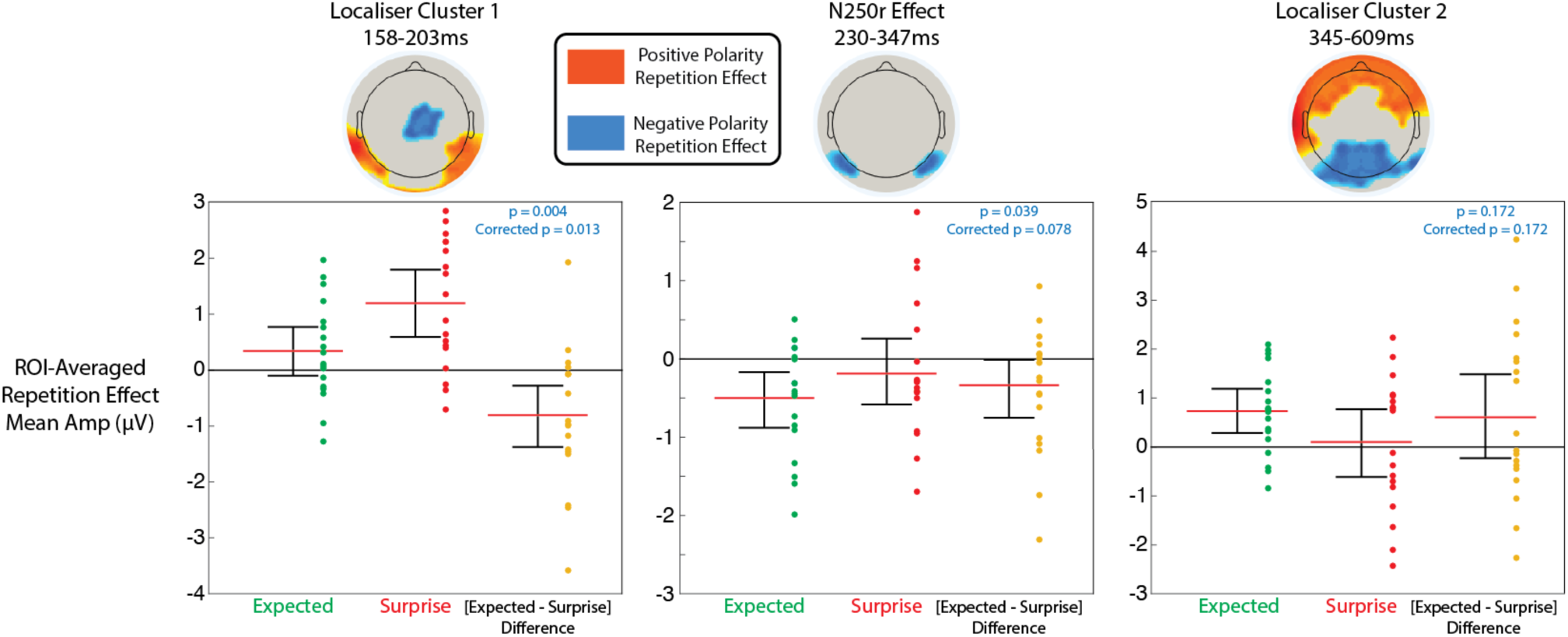
Results of ROI-based analyses of temporal expectation effects on repetition effect magnitudes. Red lines denote 20% trimmed means. Error bars denote 95% confidence intervals. Dots adjacent to error bars represent individual data points for each condition. P-values are displayed for temporal expectation by repetition interaction tests, both uncorrected for multiple comparisons and corrected using the Holm-Bonferroni method.

#### 3.3.1 Localiser cluster 1 (158-203ms)

For the combined positive-going and negative-going repetition effects in Localiser Cluster 1, repetition effects were larger for faces after surprising 500ms ISIs (trimmed mean repetition effect = 1.20*μ*V, CI = [0.59, 1.80]) compared to expected ISIs (trimmed mean repetition effect = 0.34*μ*V, CI = [-0.10, 0.79], trimmed mean [expected - surprising] difference = −0.81*μ*V, CI of the difference = [−1.38, −0.28], p = .004, Holm-Bonferroni adjusted p = .013; Figure 4 left panel).

#### 3.3.2 N250r effect (230-347ms)

During the N250r time window negative-going repetition effects were larger for faces after expected 500ms ISIs (trimmed mean repetition effect = −0.50*μ*V, CI = [−0.88, −0.16]) compared to surprising 500ms ISIs (trimmed mean repetition effect = −0.19*μ*V, CI = [−0.58, 0.27], trimmed mean [expected – surprising] difference = −0.34*μ*V, CI of the difference = [−0.74, −0.01]; Figure 4 center panel). However this effect was not statistically significant after correction for multiple comparisons (p = .039, Holm-Bonferroni adjusted p = .078).

#### 3.3.3 Localiser cluster 2 (345-609ms)

For the combined positive-going and negative-going repetition effects in Localiser Cluster 2 there were no statistically significant differences in magnitudes of repetition effects for stimuli following expected 500ms ISIs (trimmed mean repetition effect = 0.73*μ*V, CI = [0.26, 1.19]) when compared with surprising 500ms ISIs (trimmed mean repetition effect = 0.10*μ*V, CI = [-0.62, 0.78], trimmed mean [expected – surprising] difference = 0.60*μ*V, CI of the difference = [-0.25, 1.51], p = .172, Holm-Bonferroni adjusted p = .172; Figure 4 right panel).

## 4. Discussion

We comprehensively investigated the spatiotemporal dynamics of temporal expectation effects on RS in the visual system using ERPs. This permitted us to test a core hypothesis of the predictive coding model RS: that RS will be larger for stimuli with expected, rather than surprising, stimulus onset times (Auksztulewicz & Friston, 2016). We report that temporal expectation affects the magnitude of RS. Interestingly, the effects are qualitatively different across early (158-203ms) and late (230-609ms) time windows. Effects during the early time window were incompatible with the predictive coding model, whereas effects in later time windows were in line with model predictions. Our results indicate different mechanisms underlying RS for responses during early and late latencies from stimulus onset, showing that the timing of repetition effects is critical to consider when specifying underlying mechanisms.

We also replicated the complex progression of ERP face image repetition effects reported in our previous study (Feuerriegel et al., 2018a), providing further evidence for a diverse range of RS effects which extend at least until 800ms from stimulus onset.

### 4.1 Effects of Temporal Expectation on Repetition Suppression

We identified early (158-203ms) and late (230-609ms) time windows during which temporal expectations modulated RS. These temporal expectation effects differed across time windows, suggesting distinct mechanisms underlying early and late latency repetition effects.

During the early time window ERP repetition effects were larger for stimuli with surprising (rather than expected) onset times. This result is incompatible with the predictive coding model of RS (Auksztulewicz & Friston, 2016) which predicts larger repetition effects following expected onset times due to increased sensory precision. Our results are also similar to findings from studies which manipulated expectations about the identity of upcoming stimuli, which found larger BOLD repetition effects for surprising stimuli in the FFA, the occipital face area and lateral occipital cortex (e.g., Amado et al., 2016; Kovács et al., 2012; Choi et al., 2017). Interactions between expectation effects and RS have not been explicitly described within normalisation models, however one hypothesis compatible with the above findings is that RS, as described in normalisation models, reduces effects of perceptual expectation and surprise (as found in Feuerriegel et al., 2018b). This could be achieved through reductions in stimulus-driven input to excitatory neurons (e.g., via inherited adaptation: Larsson et al., 2016; Feuerriegel, 2016), which would bias competitive excitatory-inhibitory interactions across stimulus-selective cells (e.g., in models of Chelazzi et al., 1998; Kaluikhovich & Vogels, 2016). Such a mechanism could also reduce effects of subsequent inhibitory inputs to stimulus-selective neurons during recurrent network activity, (e.g., those of inhibitory interneurons: Auksztulewicz & Friston, 2016), which are associated with prediction error minimisation and perceptual expectation. This would be equivalent to RS reducing salience or sensory precision within predictive coding models (Solomon & Kohn, 2014; Feuerriegel et al., 2018b).

Later repetition effects spanning 230-609ms were larger for stimuli with expected onset times, consistent with the predictive coding model of RS. Stimuli that appeared at expected times showed repetition effect topographies consistent with repetition effects identified using the localiser dataset (Figure 3) and those in our previous study (Feuerriegel et al., 2018a). Stimuli with surprising onset times did not show such repetition effect topographies (see Figure 3C). However, the interaction effects in our analyses were not statistically significant after correction for multiple comparisons, and strong conclusions are not warranted until these effects can be replicated.

Although the interpretation of the later temporal expectation effects remains speculative, it is possible that earlier repetition effects are better described by normalisation models (designed to describe RS of stimulus-driven afferent responses) and later effects are better described by predictive coding models, which act via feedback-driven modulations of recurrent network activity in visual cortex (see Grotheer & Kovács, 2016; Grimm et al., 2016 for similar ‘hybrid’ models of RS). The possibility of multiple RS mechanisms also raises interesting questions regarding how RS of early afferent responses, as specified in normalisation models, might influence later effects of perceptual expectations within the same recurrently-connected networks, as well as across brain regions via inherited effects. Predictive coding mechanisms could potentially utilise locally-generated RS (e.g., via afterhyerpolarisation or synaptic fatigue) to implement a default expectation for recently-seen stimuli, whereby expectation and surprise effects are maximal for events that signify changes in the environment.

### 4.2 Face Image Repetition Effects

Using mass univariate analyses of ERPs we also identified a complex progression of repetition effects spanning 158-800ms from stimulus onset, with differing topographies across separate time windows (displayed in Figure 2). We replicated repetition effects reported in our previous study (Feuerriegel et al., 2018a) with regard to the polarity, latency, scalp topography and approximate magnitude of each effect.

Our results provide further evidence for multiple, distinct RS effects that occur over a wide time range following stimulus onset. Many of these effects were detected at electrodes over visual cortex, indicating that RS as measured by fMRI BOLD in the visual system reflects a mixture of early and late effects. If this is the case, it is unclear whether the effects of attention and expectation on BOLD RS (e.g., Eger et al., 2004; Summerfield et al., 2008) reflect early or late modulations of stimulus-evoked responses. Electrophysiological recordings with higher temporal precision (e.g., Todorovic & de Lange, 2012; Neumann & Schweinberger, 2009) will be needed to characterise the timing of these effects. Such timing data will be critical to identify different mechanisms underlying RS, including the relative timing of contributions by each mechanism.

The earliest (162-211ms) statistically significant repetition effect resembled reductions of N170 component amplitudes as reported in previous studies (Caharel et al., 2011; 2015). In Caharel et al. (2015) this effect was found only with identical image repetitions or small viewpoint differences between adapter and test faces, and so is likely to index repetition of low-level image features or stimulus shape. We also identified the N250r face identity repetition effect (Schweinberger et al., 2002) spanning 228-369ms from test stimulus onset. As in our previous study, we identified a mid-latency repetition effect spanning 371-619ms from stimulus onset. The latency and broad scalp topography across posterior electrodes suggests widespread secondary effects of RS on recurrent local network activity (e.g., Kaliukhovich & Vogels, 2016; Patterson, et al., 2013) or consequences of feedforward and feedback interactions across visual areas (e.g., Ewbank et al., 2011). We also replicated a late repetition effect from 720ms until the end of the 800ms epoch. Such a late effect is unlikely to result from changes in stimulus-driven afferent input, and may index feedback input into visual areas from frontal regions (Grotheer & Kovács, 2016).

One difference in results compared to our previous study is that we did not find a very early repetition effect (~100ms, around the time of the visual P1 component) to be statistically significant. However, we did find a similar topography of effects when averaging ERPs over the time window of the P1 in our experiment (Figure 2B). It is likely that our smaller sample size (n = 18 vs. n = 36) prevented us from detecting this short-duration effect using a cluster-based permutation test (see Groppe et al., 2011).

### 4.3 Caveats

Our research should be interpreted with the following points kept in mind. Firstly, it is important to note that the repetition effects reported here reflect face *image* repetition rather than face *identity* repetition. Early repetition effects, due to repetition of low-level features, may have caused later repetition effects, or enhanced their magnitudes via compounding inherited RS effects across brain regions (reviewed in Larsson et al., 2016; Feuerriegel et al., 2016). Studies aiming to isolate identity repetition effects should use different image sizes (Dzhelyova & Rossion, 2014), or different images with minimal overlap of local features (Schweinberger & Neumann, 2016).

Also, it is likely that the late (720-800ms) repetition effect described here extends beyond 800ms after stimulus onset. In both the current experiment and our previous study we could not extend our analysis epochs beyond 800ms from stimulus onset due to paradigm design (as epochs would overlap with the earliest onset times of the fixation cross in the subsequent trial). Future experiments should use longer intertrial intervals to assess whether immediate stimulus repetition effects extend even further in time than described here.

In our study we could not determine whether the modulations of repetition effects in our experiment are due to temporal attention or expectation. Previous studies of temporal attention have also manipulated expectations regarding the temporal onset of task-relevant stimuli (e.g. Miniussi et al., 1999; Griffin et al., 2002) and so far not many experiments have separately manipulated temporal expectation and attention (but see Schwartze et al., 2011, 2013; Paris et al., 2015). While either effect (temporal attention or temporal expectation) yields the same hypotheses according to the predictive coding model of RS, the distinction between expectation and attention will likely be important when developing mechanistic models of RS effects within local neural networks. Future work that orthogonally manipulates temporal attention and expectation will be able to distinguish between these effects.

### 4.4 Conclusions

The research reported here shows that temporal expectations modulate RS, and that modulatory effects differ across early and late phases of stimulus-evoked responses. These differing effects of temporal expectations indicate that multiple, different mechanisms likely underlie RS in the visual system, depending on the timing and anatomical location of each effect. Our results support the development of hybrid RS models, which specify multiple, potentially interacting effects of RS, expectation and attention within sensory systems (e.g., Grotheer & Kovács, 2016; Grimm et al., 2016).

## Acknowledgements

We thank Dilushi Chandrakumar for her assistance with preparation of stimuli, and Rik Henson and Jason Mattingley for their feedback on an earlier version of this manuscript. Daniel Feuerriegel was supported by an Australian Government Research Training Program Scholarship. Hannah Keage is supported by a NHMRC Dementia Research Leadership Fellowship (GNT1135676). Funding sources had no role in study design, data collection, analysis or interpretation of results.

